# Transplanted cells are essential for the induction but not the expression of cortical plasticity

**DOI:** 10.1101/644377

**Authors:** Mahmood S. Hoseini, Benjamin Rakela, Quetzal Flores-Ramirez, Andrea R. Hasenstaub, Arturo Alvarez-Buylla, Michael P. Stryker

## Abstract

Transplantation of even a small number of embryonic inhibitory neurons from the medial ganglionic eminence (MGE) into postnatal visual cortex makes it lose responsiveness to an eye deprived of vision when the transplanted neurons reach the age of the normal critical period of activity-dependent ocular dominance (OD) plasticity. The transplant might induce OD plasticity in the host circuitry or might instead construct a parallel circuit of its own to suppress cortical responses to the deprived-eye. We transplanted MGE neurons expressing archaerhodopsin, closed one eyelid for 4-5 days, and, as expected, observed transplant-induced OD plasticity. This plasticity was evident even when the activity of the transplanted cells was suppressed optogenetically, demonstrating that the plasticity was produced by changes in the host visual cortex.

**Significance Statement:** Interneuron transplantation into mouse V1 creates a window of heightened plasticity which is quantitatively and qualitatively similar to the normal critical period, i.e. short-term occlusion of either eye markedly changes ocular dominance. The underlying mechanism of this process is not known. Transplanted interneurons might either form a separate circuit to maintain the ocular dominance shift or might instead trigger changes in the host circuity. We designed experiments to distinguish the two hypotheses. Our findings suggest that while inhibition produced by the transplanted cells triggers this form of plasticity, the host circuity is entirely responsible for maintaining the ocular dominance shift.

**One Sentence Summary:** Neuronal transplants do not just grow and connect—they induce plasticity in the adult brain.

## Introduction

Normal brain development is marked by temporally restricted windows of heightened experience-dependent plasticity known as critical periods (CPs). During a CP in primary visual cortex (V1)—postnatal day (P) 25-35 in mouse—changes in neuronal circuitry facilitate the matching of left eye and right eye receptive fields in V1 binocular neurons (reviewed in (Espinosa and Stryker, 2012)). Occlusion of vision of either eye over this time, referred to as monocular deprivation (MD), prevents binocular matching and results in structural and functional changes that reduce neural responses to the deprived eye and increase them to the eye that remains open (Wiesel and Hubel, 1963; Wang et al., 2010).

Local inhibitory interneurons in V1 play a crucial role in opening the CP for ocular dominance (OD) plasticity, which begins ~35 days after these neurons are generated in the medial ganglionic eminence (MGE) (Hensch et al., 1998; Hanover et al., 1999; Pizzorusso, 2002). Once the CP has closed, V1 circuits are thought to remain stable throughout life. However, transplantation of newly generated parvalbumin or somatostatin interneurons from the MGE of a donor embryo into the V1 of a postnatal host opens a second brief CP ~35 days after transplantation (DAT) (Southwell et al., 2010; Tang et al., 2014; Davis et al., 2015). Similar transplantation has also shown promise in improving visual acuity in adult mice after long-term MD (Davis et al., 2015).

To dissect the precise contribution of inhibitory interneurons in OD plasticity, it is essential to draw a distinction between induction and expression of the OD shift. Induction of OD shift consist of a set of mechanisms (e.g. enhanced inhibition) that facilitate the occurrence and magnitude of the OD shift (Feldman, 2000). On the other hand, expression refers to the changes in cortical circuit that reduce the response to the deprived eye persistently (Gandhi et al., 2008). While inhibitory neuron activity is crucial in inducing the normal CP (Hensch et al., 1998; Kuhlman et al., 2013), the expression of the plasticity following MD no longer depends on inhibition (Saiepour et al., 2015). Here, we investigated whether the MGE transplants produce a second CP of OD plasticity by stimulating changes in host circuitry or, alternatively, by constructing a separate parallel circuit within the host tissue to inhibit deprived-eye responses and disinhibit non-deprived-eye responses. We transplanted inhibitory interneurons expressing either archaerhodopsin-3 or channelrhodopsin-2 that allowed us to silence or activate the transplanted interneurons after plasticity is induced by 4-5 days of MD during the second CP. Our results indicate that an OD shift induced by brief MD persists under conditions of either reduced or enhanced transplant-derived inhibition. While the manipulating of the activity of transplanted interneurons after plasticity has been induced alters the ODs of some individual neurons, it does not change the overall OD at the population level. Our findings reveal that transplanted interneurons are not responsible for the expression of OD plasticity but rather facilitate its induction in the host circuitry.

## Materials and Methods

### Experimental animals

All protocols and procedures followed the guidelines of the Animal Care and Use Program at the University of California, San Francisco under IACUC Protocol AN143347. All mice were housed in standard conditions on a 12h dark/light cycle. Experiments were performed during the dark, more active phase of the cycle. Ai40 (JAX Stock No: 021188), Ai32 (012569), Gad2-Cre (010802), and wild-type C57BL/6J (000664) lines were purchased from The Jackson Laboratory and maintained in our housing facility.

### Cell dissection and transplantation

The ventricular and subventricular zones of the MGE were dissected from E13.5 donor embryos as in Vogt et al. (Vogt et al., 2015). The tissue was dissociated by repeated pipetting in Leibovitz’s L-15 medium containing 100U/mL DNase I (Sigma Aldrich, D5025-15KU). Dissociated cells were concentrated via centrifugation (800 rcf for 4 minutes). P5-8 recipients were anesthetized via hypothermia until pedal reflex disappeared and then placed on a stereotaxic platform for injection of concentrated cells (250-350 cells/nL, 150 nL per injection for 120-200K cells total injected) through a beveled Drummond glass micropipette (Drummond Scientific) positioned at 45 degrees from vertical. Three injections were placed into the caudal left cortex at 1.5 mm anterior from lambda, 3.5 mm lateral from midline and 1.5 mm anterior, 1 mm lateral, and 3 mm anterior, 1.25 mm lateral. Injections were made at 0.5-1.0 mm deep from the skin surface. After injection, recipients were placed on a heating pad until warm and active at which time they were returned to their mothers until weaning (P28).

### Headplate surgery and alert recording apparatus

To allow for stable recordings, the animal’s head was fixed to a rigid crossbar above the floating ball. This was achieved by attaching a titanium headplate (circular center with a 5 mm central opening) to the skull a week before the recording [28 days after transplantation (DAT)].

For headplate attachment, animals were anesthetized with isoflurane in oxygen (3% induction, 1.5% maintenance) and given a subcutaneous injection of Meloxicam (10 mg/kg) to reduce inflammation, a subcutaneous injection (0.05 ml) of lidocaine above the skull as a local anesthetic, and a subcutaneous injection of 0.2 ml of saline to prevent dehydration. After a scalp incision, the fascia was cleared from the surface of the skull and the titanium headplate was then attached with Metabond (Parkell Co.), covering the entire skull, except for the region in the center of the headplate, which was covered with a 0.2% benzethonium chloride solution (New-Skin Liquid Bandage) to protect the skull. After recovery, the animal was allowed to habituate to the recording setup by spending about one hour on the floating ball over the course of 2–3 days, during which time the animal was allowed to run freely. The polystyrene ball was constructed using two hollow 200-mm-diameter halves (Graham Sweet Studios) placed on a shallow polystyrene bowl (250 mm in diameter, 25 mm thick) with a single air inlet at the bottom. Two optical USB mice, placed 1 mm away from the edge of the ball, were used to sense rotation of the floating ball and transmit signals to our data analysis system using custom driver software. These measurements are used to divide data into still and running trials and analyze them separately.

### Intrinsic signal imaging

Five days prior to recording, mice were anesthetized with 0.7% isoflurane and a single i.p. injection of 2-5 mg/kg chlorprothixene. The headplate was secured to a stereotaxic frame, filled with agarose, and topped with a coverslip to provide a flat imaging surface. Animals were maintained at 37.5°C, measured using a rectal thermometer, and eyes were protected with silicone oil. Intrinsic signal images were acquired at 30 frames per second using a Dalsa 1M30 CCD camera with a red interference filter. The camera was focused 550μm from the cortical surface and collected 610 nm light that was reflected off the visual cortex through the skull.

Visual response amplitude was defined as the average signal amplitude for each pixel and ocular dominance index was computed as (C−I)/(C+I) where C is the response to contralateral visual stimulation and I is the response to ipsilateral visual stimulation (Cang et al., 2005).

### Monocular deprivation and microelectrode recording

Four or five days prior to extracellular recording, monocular deprivation was induced by suturing the right eyelid shut and was checked daily to confirm eyelid closure (**Fig. 1A**).

**Fig. 1.**
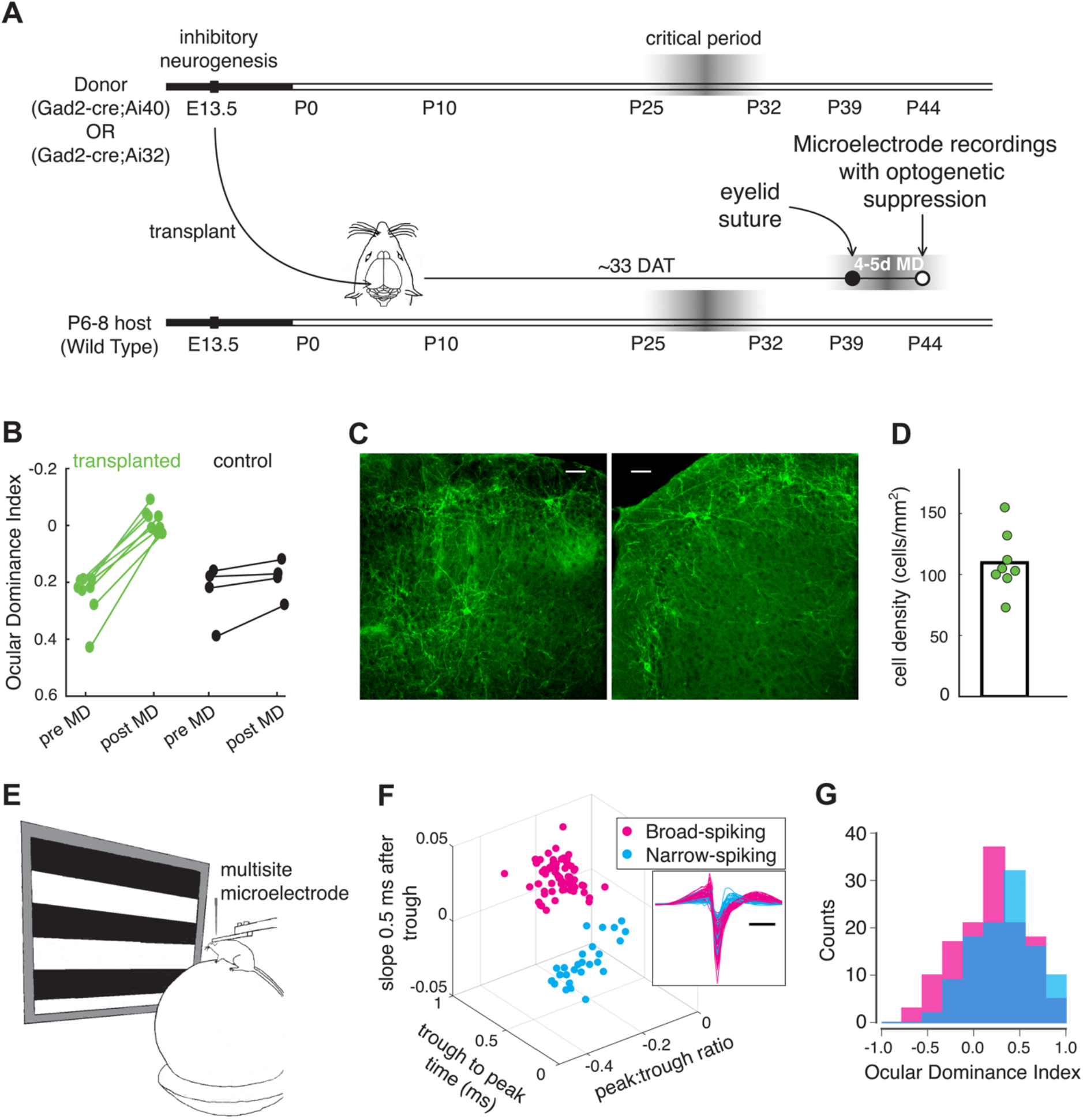
Transplanting medial ganglionic eminence (MGE)-derived interneurons induces a second critical period (CP). (A) Cortical GABAergic interneurons are produced from E13.5 embryos of crossing Gad2-cre mouse line with either Ai40 (RCL-ArchT-EGFP) or Ai32 (RCL-ChR2-EYFP). Mouse visual cortex shows one week of heightened plasticity from postnatal day (P) 25 to 32, when inhibitory neurons are 33 to 35 days old. Similarly, inhibitory neuron precursors transplanted from E13.5 donor embryos into P6 hosts create a second critical period from P38-45. Following 4-5 days of monocular deprivation (MD) visual responses were recorded using multisite microelectrodes in the presence of optogenetic manipulations. (B) Ocular dominance (OD) index shows significant plasticity following 4-5 days of MD in the transplant-recipient mice relative to the control animals as measured by optical imaging of intrinsic signal (transplanted: mean±SEM=0.24±0.03 to −0.01±0.01, Mann-Whitney test, U=0, p=0.0005, n=8 mice; control: 0.24±0.05 to 0.18±0.03, U=6, p=0.33, n=4 mice). (C) Coronal brain sections, harvested after microelectrode recordings, at P46 show transplanted cells engrafted and functionally integrated in the binocular zone of the visual cortex. (D) The overall density of transplanted cells in the host primary visual cortex at P46 (109±9, transplant-derived neurons per mm^2^; n = 8 mice). (E) Recordings were acquired from the binocular visual cortex of freely-moving mice while presenting drifting gratings (12 orientations, 1.5 s duration, and 1.5 s inter-trial interval). (F) Using the three parameters calculated from average waveforms, cells were classified into narrow- (inhibitory, cyan) or broad- (excitatory, magenta) spiking (height of the positive peak relative to the negative trough: * 0.209±0.008, −0.287±0.018, slope of the waveform 0.5 ms after the negative trough: 0.019±0.001, −0.011±0.001, the time from the negative trough to the peak: 0.748±0.010, 0.313±0.016 ms, excitatory and inhibitory cells respectively). Shown in the subplot is the average spike waveforms for all units, aligned to minimum, demonstrating excitatory (magenta; 61) and inhibitory (cyan; 26) cells. (G) The histogram of OD index across excitatory and inhibitory populations in control mice without MD (average ODI, 0.18±0.03, n=132 excitatory cells, 3 mice; 0.33±0.03, n=108 inhibitory cells; p=0.002, Wilcoxon rank-sum test)

Extracellular recording was carried out in awake, head-fixed mice that were free to run on the floating polystyrene ball (**Fig. 1E**). On the day of recording, the animal was anesthetized with isoflurane (3% induction, 1.5% maintenance) and sutures were removed to open the eyelid. The liquid bandage was removed from the skull, and a craniotomy of about 1-2 mm in diameter was made above the binocular zone of V1 (identified by intrinsic signal imaging). This small opening was large enough to allow for insertion of a 1.1-mm-long double-shank 128-channel electrode (Du et al., 2011), fabricated by the Masmanidis laboratory (University of California—Los Angeles) and assembled by the Litke laboratory (University of California—Santa Cruz). After animals were recovered, the electrode was placed at an angle of 20-40° to the normal of the cortical surface and inserted to a depth of 500-1000 μm. An optical fiber (400 μm diameter) coupled to a light source (green LED for archaerhodopsin, peak 595 nm; blue LED for channelrhodopsin, peak 473 nm) was positioned above the cranial window, centered on the binocular region, to illuminate V1. Recordings were started an hour after implantation. For each animal, the electrode was inserted not more than twice.

### Visual stimuli

Stimuli were displayed on an LCD monitor (Dell, 30×40 cm, 60 Hz refresh rate, 32 cd/m^2^ mean luminance) placed 25 cm from the mouse (−20° to +40° elevation) without gamma correction. Drifting sinusoidal gratings at 12 evenly spaced directions (40 repetitions, 1.5 s duration, 0.04 cycles per degree, and 1 Hz temporal frequency) were generated and presented in random sequence using the MATLAB Psychophysics Toolbox (Brainard, 1997; Kleiner et al., 2007) followed by 1.5-second blank period of uniform 50% gray.

### Data acquisition

Movement signals from the optical mice were acquired in an event-driven mode at up to 300 Hz, and integrated at 100 ms intervals and then converted to the net physical displacement of the top surface of the ball. A mouse was said to be running on a single trial if its average speed for the first 500 ms of the trial fell above a threshold, found individually for each mouse (1–3 cm/s), depending on the noise levels of the mouse tracker. Data acquisition was performed using an Intan Technologies RHD2000-Series Amplifier Evaluation System, sampled at 20 kHz; recording was triggered by a TTL pulse at the moment visual stimulation began.

### Single-neuron analysis

Single units were identified using MountainSort (Chung et al., 2017), which allows for fully automated spike sorting of the data acquired using 128-site electrodes and runs at approximately 2x real time. Manual curation after a run on one hour of data takes an additional half hour, typically yielding 90 (range 50-130) isolated single units. Average waveforms of isolated single units were used to calculate three parameters by which cells were classified into narrow- or broad-spiking (Niell and Stryker, 2008). The parameters are: the height of the positive peak relative to the negative trough, the slope of the waveform 0.5 ms after the negative trough, and the time from the negative trough to the peak (**Fig. 1F**). Cells were discarded under one of the following criteria, (i) average firing rate varies more than 50% across trials, (ii) not responding above baseline to either eye, (iii) substantially different tuning curves in response to the stimulation of the two eyes. Overall ~40% (range 16-67) of the cells passed the neuronal exclusion criteria.

### Immunostaining

To determine whether transplanted cells were in the binocular zone of V1, animals were perfused transcardially after microelectrode recording first with ice-cold PBS and then with a solution of 4% paraformaldehyde (PFA) prepared in PBS. The brains were harvested and postfixed overnight in a solution of 4% PFA and subsequently cryoprotected by immersion in a PBS solution containing 30% sucrose. The brains were sectioned coronally (40 μm thickness) using a sliding microtome (Leica; Physitemp Instruments). Free-floating sections were permeabilized in 0.25% Triton X-100/ PBS solution, blocked for an hour at room temperature in a blocking solution containing tris-buffered saline, 10% normal goat serum, and 0.1% Triton X-100. They were then incubated overnight at 4C with 1:500 diluted chicken anti-GFP (Abcam ab13970) primary antibody. After washing in PBS five times for 10 minutes each, sections were incubated in blocking solution with goat anti-chicken secondary antibodies (Thermo Fisher, Alexa Fluor 488, A-11039) at 1:1000 for 1 hour at room temperature. They were then washed three times in PBS for 10 minutes each, mounted on glass slides using mounting media, and coverslipped.

### Cell counting

Cell density was defined as the number of fluorescent cells within the binocular visual cortex divided by the total area. Binocular visual cortex was identified functionally during intrinsic signal imaging sessions. Fluorescent images of the binocular visual cortex were imaged using Nikon Eclipse 90i inverted microscope with a Nikon 4x objective.

### Experimental Design and Statistical analysis

The experiments reported here were designed to determine (1) whether optogenetic suppression/activation of the transplanted interneurons alters network activity and/or ocular dominance, and (2) whether delivering the same optogenetic light affects network activity and/or ocular dominance in the control mice. For experiment (1), we recorded 10 mice (6 females and 4 males) that were transplanted with interneurons expressing ArchT (suppression) and 5 mice (2 females and 3 males) that were transplanted with interneurons expressing ChR2 (activation). All data are illustrated in Figs. 1, 2, 4 and tables 1 and 2. For experiment (2), we recorded 4 mice (2 females and 2 males) that did not receive any transplant. All data are shown in the scatter plots of Figs. 3 and 4. The main source of variability for all of these experiments is the individual neurons. Statistical significance was done in MATLAB and Python using a Mann-Whitney U, two-sample Kolmogorov-Smirnov, Wilcoxon rank-sum, Wilcoxon signed-rank test. Correction for multiple comparisons were done using the Benjamini-Hochberg procedure with a false discovery rate of 10%. All data are presented as mean ± SEM unless stated otherwise.

**Fig. 2.**
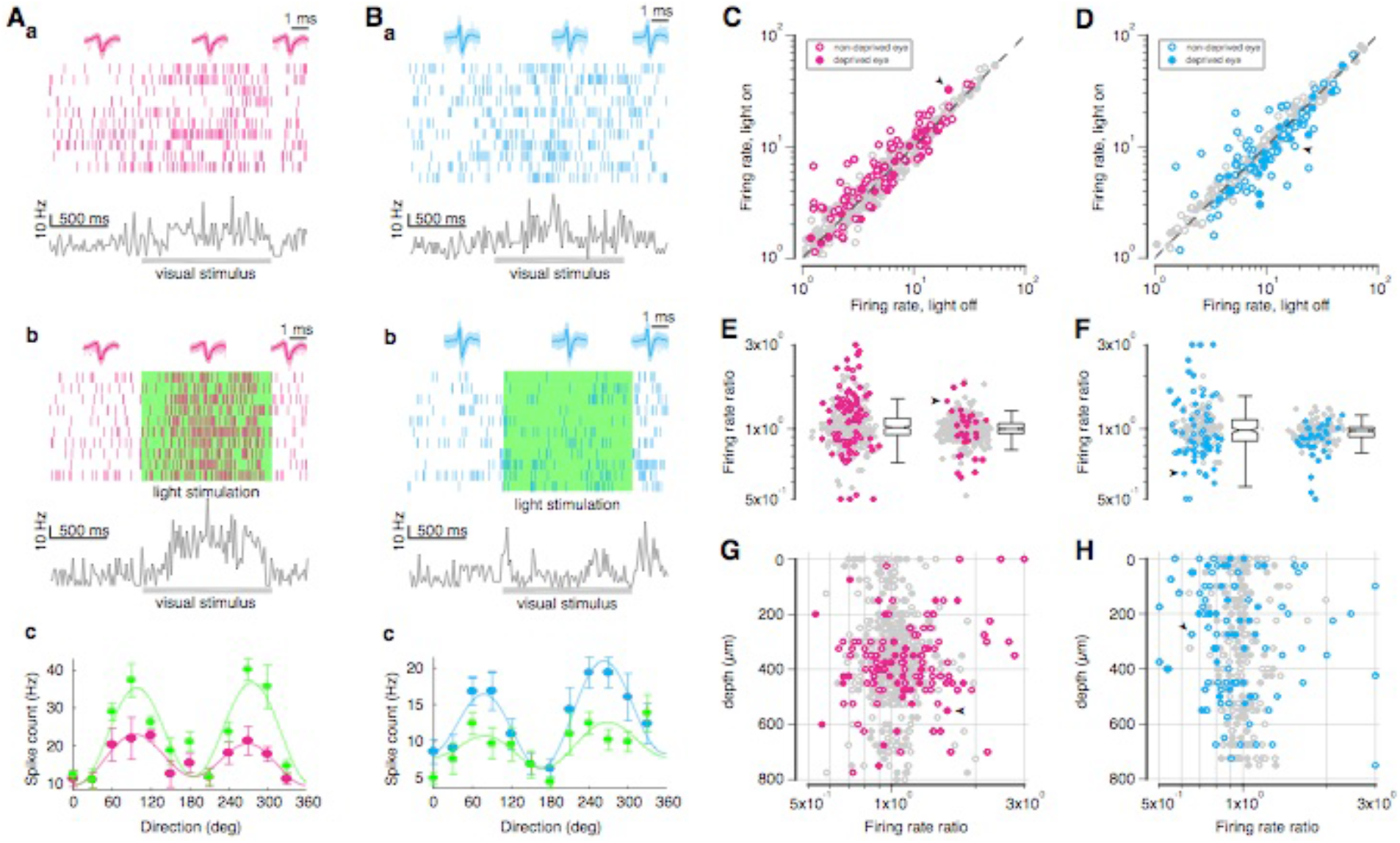
Suppressing MGE-derived transplanted interneurons modulates network activity across different populations and cortical depths. (A) An example excitatory cell that shows an increase in firing rate with optogenetic suppression of transplanted cells. a. Spike raster for a representative excitatory neuron across trials with its instantaneous firing rate shown at the bottom. Shaded area shows the duration of the stimulus. Shown at the top are the individual (low opacity) overlaid with average (high opacity) waveforms for epochs before, during, and after stimulus presentation. b. Same as in a with optogenetic silencing. Green shaded shows the duration of optogenetic suppression (green light, 532 nm). c. Tuning curves for evoked responses in control (magenta) and optogenetic suppression (green) conditions. Solid curves are fit to a sum of two sinusoids. (B) Same as in A for an example inhibitory neuron suppressed by optogenetic suppression. (C) Evoked mean firing rate, averaged over all visual stimuli for a 1-second window following stimulus onset, of excitatory cells with optogenetic suppression (light on) against control (light off) condition in response to stimulation of both eyes (slope=1.03, r=0.98). Colored dots represent cells that have significant firing rate modulation. Of 273 excitatory cells, 82 and 29 cells are significantly modulated in response to ipsi- (open circles) and contra-lateral (solid circles) eyes (Benjamini-Hochberg adjusted p=0.029), 97 cells show significant modulation in response to either eye stimulation (10 mice). Gray dashed line indicates unity. Arrowhead indicates the example cell shown in A. (D) Same as in C for inhibitory cells (slope=0.97, r=0.97). Colored dots represent cells that have significant firing rate modulation. Of 136 inhibitory cells, 66 and 32 cells are significantly modulated in response to ipsi- (open circles) and contra-lateral (solid circles) eyes (Benjamini-Hochberg adjusted p=0.055), 79 cells show significant modulation in response to either eye stimulation. Arrowhead indicates the example cell shown in B. Colored dots represent cells that have significant firing rate modulation. (E) The ratio of light on to light off average firing rates of excitatory neurons for non-deprived- (left, 46 and 36 cells positively and negatively modulated, respectively; 1.073±0.001) and deprived- (right, 14 and 15 cells positively and negatively modulated, respectively; 1.01±0.001) eye responses. (F) Same as in E for inhibitory cells (left, 24 and 42; right, 9 and 23 cells positively and negatively modulated, respectively; 1.073±0.003, 0.964±0.001). (G) Change in the ratio of mean firing rates against cortical depth, measured from the top channel in the brain, for excitatory cells (ρ=0.09, p=0.08, Spearman rank-order correlation). (H) Same as in G for inhibitory cells (ρ=0.01, p=0.89).

**Fig. 3.**
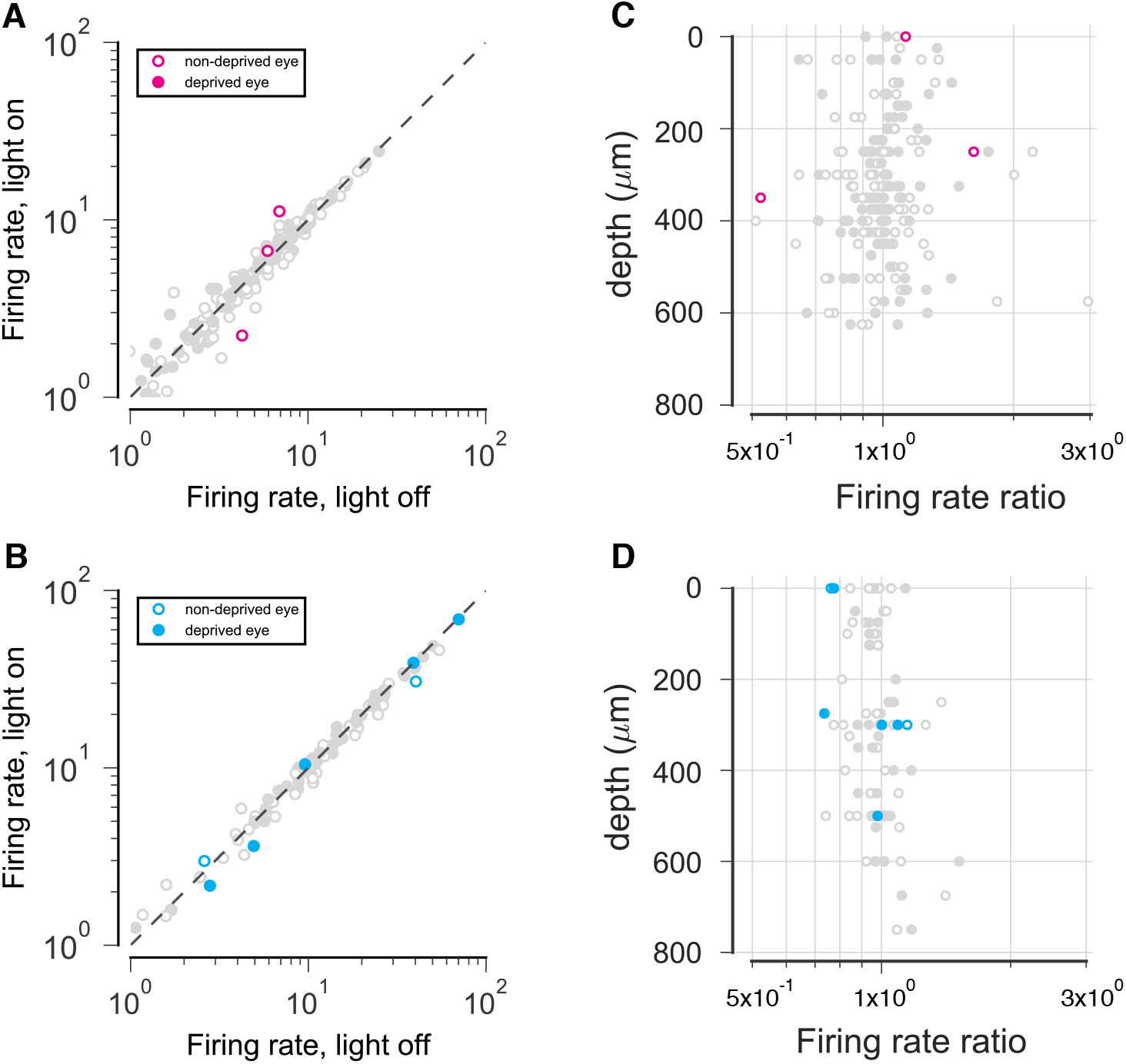
Optogenetic stimulation does not change network activity in the control mice. (A) Evoked mean firing rate, averaged over 1-second window of all visual stimuli, of excitatory cells with optogenetic stimulation (light on; green light, 532 nm) against control (light off) condition in response to stimulation of both eye (solid circles, deprived eye; open circles, non-deprived eye). Colored dots represent cells that show significant change in in their firing rate. Of 105 excitatory cells, 3 and 0 cells are significantly modulated in response to ipsi- (open circles) and contra-lateral (solid circles) eyes (Benjamini-Hochberg adjusted p=0.003; 4 mice). Gray dashed line indicates unity. (B) Same as in A for inhibitory cells. Of 55 inhibitory cells, 2 and 5 cells are significantly modulated in response to ipsi- (open circles) and contra-lateral (solid circles) eyes (Benjamini-Hochberg adjusted p=0.020). (C) Change in the ratio of firing rates against cortical depth for excitatory cells (ρ=−0.11, p=0.26, Spearman rank-order correlation). (D) same as in C for inhibitory cells (ρ=0.10, p=0.43).

**Fig. 4.**
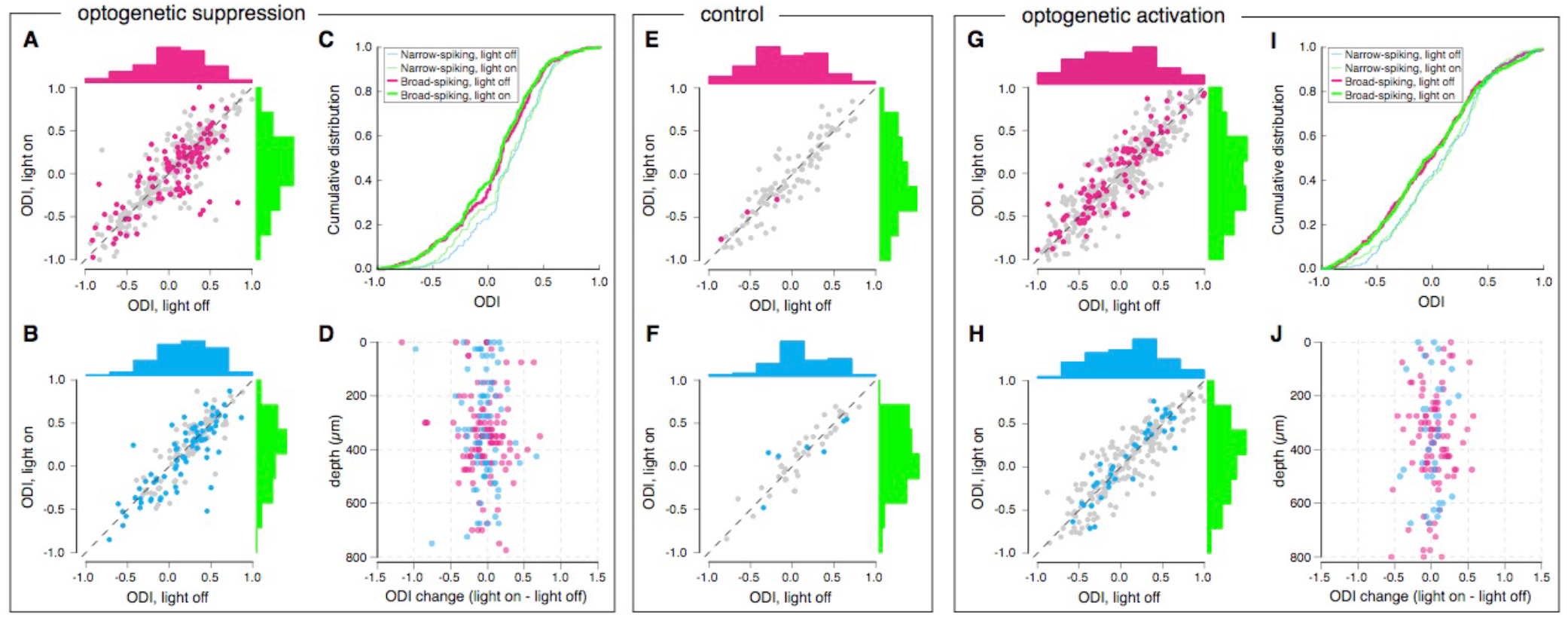
Excitatory and inhibitory population-level OD indices (ODI) are not altered by suppressing/activating transplanted interneurons. (A) Scatterplots of ODIs of excitatory cells under light on versus light off conditions. Distribution of ODIs is shown as histogram on the top (light off, magenta) and right (light on, green) (light off to light on: 0.067±0.019 to 0.046±0.019, p=0.12, Wilcoxon signed-rank test; n=273 cells, 10 mice). (B) Same as in A for inhibitory cells (light off to light on: 0.189±0.024 to 0.166±0.027, p=0.12; n=136 cells, 10 mice). (C) Cumulative distribution for the light on (green; thick curve, excitatory; thin curve, inhibitory) and light off (magenta, excitatory; cyan, inhibitory). (D) Change in the ODI against cortical depth for excitatory cells (magenta) and inhibitory cells (cyan) (excitatory, ρ=0.05, p=0.26; inhibitory, ρ=0.09, p=0.23; Spearman rank-order correlation). (E) Same as in A for control mice (light off to light on: −0.062±0.038 to −0.071±0.041, p=0.43, Wilcoxon signed-rank test; n=105 cells, 4 mice). (F) Same as in B for control mice (light off to light on: 0.098±0.046 to 0.112±0.047, p=0.60; n=55 cells). (G) Same as in A but in the presence of optogenetic activation (light off to light on: −0.031±0.023 to −0.026±0.023, p=0.76, Wilcoxon signed-rank test; n=405 cells, 5 mice). (H) Same in G for inhibitory cells (light off to light on: 0.088±0.025 to 0.081±0.026, p=0.57; n=244). (I) Same as in C in the presence of optogenetic activation (excitatory, p=0.97; inhibitory, p=0.92; two-sample Kolmogorov-Smirnov test). (J) Same as in D in the presence of optogenetic activation (excitatory, ρ=−0.08, p=0.11; inhibitory, ρ=0.05, p=0.43; Spearman rank-order correlation).

**Table 1.** Number of cells significantly modulated by optogenetic manipulation of transplanted cells in response to ipsilateral eye.

| Mouse no.<br>(sex) | M #1<br>(M) | M #2<br>(F) | M #3<br>(F) | M #4<br>(F) | M #5<br>(F) | M #6<br>(M) | M #7<br>(M) | M #8<br>(F) | M #9<br>(F) | M #10<br>(M) | total |
| --- | --- | --- | --- | --- | --- | --- | --- | --- | --- | --- | --- |
| Excitatory | 3/12 | 12/31 | 15/42 | 7/37 | 10/27 | 6/17 | 6/42 | 6/42 | 8/20 | 9/21 | 82/273 |
| Inhibitory | 2/4 | 13/20 | 12/25 | 9/17 | 5/15 | 4/12 | 10/21 | 4/8 | 4/8 | 3/6 | 66/136 |

**Table 2.** Number of cells significantly modulated by optogenetic manipulation of transplanted cells in response to contralateral eye.

| Mouse no.<br>(sex) | M #1<br>(M) | M #2<br>(F) | M #3<br>(F) | M #4<br>(F) | M #5<br>(F) | M #6<br>(M) | M #7<br>(M) | M #8<br>(F) | M #9<br>(F) | M #10<br>(M) | total |
| --- | --- | --- | --- | --- | --- | --- | --- | --- | --- | --- | --- |
| Excitatory | 2/12 | 2/31 | 2/42 | 4/37 | 4/27 | 1/17 | 4/42 | 1/24 | 5/20 | 4/21 | 29/273 |
| Inhibitory | 1/4 | 4/20 | 3/25 | 3/17 | 5/15 | 3/12 | 5/21 | 2/8 | 3/8 | 3/6 | 32/136 |

## Results

Transplantation of MGE-derived interneurons into the visual cortex of adult mice induces a second CP of ocular dominance plasticity (Southwell et al., 2010). It is unknown what circuit changes are responsible for induction and maintenance of this form of OD plasticity. More specifically, is transplant-induced OD plasticity a consequence of changes in host circuitry like those in the normal CP, or is it instead the product of a parallel circuit constructed by the transplanted neurons themselves? To discriminate between the two alternative hypotheses, we transplanted MGE neurons expressing either archaerhodopsin-3 (RCL-ArchT-EGFP) (Han et al., 2011) or channelrhodopsin-2 (RCL-ChR2-EYFP) (Madisen et al., 2012) to enable us to silence or activate them after plasticity was induced by 4-5 days of MD during the a second CP ~33 DAT (**Fig. 1A**).

Embryonic MGE-derived cells expressing excitatory or inhibitory opsins at 13.5 days of gestation were transplanted into the primary visual cortex of wild type recipients, and they dispersed, engrafted, and functionally integrated into the host cortex. They also induced a second CP (33 DAT) (**Figure 1B**; transplanted: mean±SEM=0.24±0.03 to −0.01±0.01, Mann-Whitney test, U=0, p=0.0005, n=8 mice; control: 0.24±0.05 to 0.18±0.03, U=6, p=0.33, n=4 mice), as previous studies have shown even when the number of transplanted interneurons is very low (Tang et al., 2014; Larimer et al., 2016) (typically <10% of the endogenous cells, **Fig. 1C, D**). The transplant-induced CP is similar in duration to the normal CP (Southwell et al., 2010; Tang et al., 2014).

After 4-5 days of MD during the transplant-induced CP, each eye’s responses to visual stimuli were recorded using multisite microelectrodes (**Fig. 1E**) to evaluate ocular dominance with and without optogenetic perturbation of the activity of the transplanted neurons on interleaved stimulus trials. Average spike waveforms were used to calculate three parameters by which cells could be classified unambiguously into narrow- or broad-spiking (Niell and Stryker, 2008) (**Fig. 1F**). Narrow-spiking cells are almost exclusively fast-spiking interneurons, while broad-spiking cells are ~90% excitatory and 10% inhibitory cells (Connors and Kriegstein, 1986; Barthó et al., 2004; Atencio and Schreiner, 2008). Hereafter, for clarity, we refer to the narrow- and broad-spiking units as inhibitory and excitatory neurons respectively. Recordings from normally reared mice reveal that inhibitory neurons respond more strongly to the contralateral eye and have corresponding larger ODI (**Fig. 1G**; average ODI, 0.18±0.03, n=132 excitatory cells, 3 mice; 0.33±0.03, n=108 inhibitory cells; p=0.002, Wilcoxon rank-sum test).

First, we verified that the optogenetic suppression of the transplanted interneurons was effective in altering network activity. Since the number of transplanted interneurons is very low and they disperse widely throughout the recipient tissue, the probability of recording from them is low. Suppressing the activity of transplanted cells after they induce plasticity (37-38 DAT), did however modulate the activity in other cells of the host circuit (**Fig. 2A, B**). The modulation of both excitatory and inhibitory populations was in both directions: the visual responses of some cells increased and others decreased, while the majority of the cells did not show any significant changes in their firing rate (**Fig. 2C, D**; excitatory: 82 and 29 out of 273 significantly modulated in response to ipsi- (open circles) and contra-lateral (solid circles) eyes, Benjamini-Hochberg adjusted p=0.029; inhibitory: 66 and 32 out of 136, p=0.055; 10 mice). Significant increases were more common in both cell population. The seemingly contradictory increases in inhibitory- and decreases in excitatory-cell firing rates are, however, expected in a balanced inhibition-stabilized network and only a minority of cells are expected to show a detectable change in activity under perturbation (see Fig. 10 in (Sadeh et al., 2017)). While reducing transplant derived inhibition causes an increase in some excitatory cells’ firing rate, this enhanced excitation might activate other inhibitory cells that in turn feedback to suppress other excitatory cells (Moore et al., 2018). **Figures 2E** and **F** show that the ratio of firing rates during optogenetic suppression (light on) to control (light off) non-deprived-eye responses are more strongly modulated in both populations. In addition, optogenetic suppression caused a greater increase in non-deprived-eye than deprived-eye responses among excitatory cells and a larger decrease in deprived-eye responses among inhibitory cells (**Fig. 2E, F**). Significant changes in the activity of both types of cells were observed across all cortical layers and in all 10 mice studied (**Fig. 2G, H**; **Tables 1** and **2**).

To control for the non-specific light effects on the brain, we performed similar experiments on the control mice lacking opsin-containing interneurons. As expected, delivering the same optogenetic stimulus light to V1 in control mice did not change network activity (**Fig. 3**).Only 3 of the 105 excitatory cells were significantly modulated in response to ipsilateral eye (Benjamini-Hochberg adjusted p=0.003; 4 mice), while 2 and 5 of 55 inhibitory cells displayed firing rate change in response to ipsi- and contralateral eyes respectively (adjusted p=0.020).

Microelectrode recordings of the host cells confirmed the OD shift disclosed by intrinsic signal imaging (**Fig. 4A**, excitatory: 0.067±0.019; **Fig. 4B**, inhibitory: 0.189±0.024). Having verified that optogenetic stimulation was effective in suppressing the activity of transplanted cells, we then asked whether the plasticity produced by MD persisted when transplanted-cell activity was suppressed. We found that suppressing transplanted cells did not cause a consistent change in the OD index of excitatory (**Fig. 4A**; optogenetic excitation light off to light on: 0.067±0.019 to 0.046±0.019, p=0.12, Wilcoxon signed-rank test; n=273 cells, 10 mice) or inhibitory (**Fig. 4B**; light off to light on: 0.189±0.024 to 0.166±0.027, p=0.12; n=136 cells, 10 mice) cells. Moreover, we did not observe a consistent change in the ODI of the cells that were significantly affected by the optogenetic stimulus (p=0.3, excitatory; p=0.59, inhibitory; Wilcoxon rank-sum test), consistent with previous reports in the normal CP (Khibnik et al., 2010; Saiepour et al., 2015).

The difference of OD distributions between excitatory and inhibitory cells raises the possibility that they may show different plasticity. Cumulative distributions of OD index indicate that inhibitory cells still have higher OD indices than excitatory cells (**Fig. 4C**; light off, p=0.002; light on, p=0.005; two-sample Kolmogorov-Smirnov test), showing that the two populations are more or less similarly affected by MD. Consistent with the fact that the optogenetic stimulation affects activity across all cortical depth (**Fig. 2G, H**), no consistent relationship was observed between changes in OD index and cortical depth (**Fig. 4D**; excitatory, ρ=0.05, p=0.26; inhibitory, ρ=0.09, p=0.23; Spearman rank-order correlation). Again, delivering the same optogenetic stimulus light to V1 in control mice did not change OD index (**Fig. 4E, F**;light off to light on: −0.062±0.038 to −0.071±0.041, n=105 excitatory; 0.098±0.046 to 0.112±0.047, n=55 inhibitory cells from 4 mice).

An alternative approach to determine whether transplantation specifically inhibits deprived-eye responses after MD is to activate the transplanted cells. Since optogenetic activation and inactivation of interneurons need not produce symmetric effects (Phillips and Hasenstaub, 2016; Moore et al., 2018), the outcome of this experiment is not trivial. To activate transplanted cells optogenetically, donor embryos expressing the gene for channelrhodopsin-2 in all of the inhibitory neurons were generated by crossing Gad2-Cre mouse line with Ai32 (RCL-ChR2-EYFP) line. After transplantation at P6 and 4-5 days of MD 35 days thereafter, similar recordings were performed (**Fig. 1A**). As expected, activating transplanted interneurons altered activity in both cell types and across cortical depths (data not shown). However, it did not consistently favor one eye over the other, and OD index did not significantly change in either excitatory or inhibitory cells (**Fig. 3G-J**; excitatory, ODI change, 0.005±0.010, p=0.76, n=405 cells from 5 mice; inhibitory, −0.007±0.013, p=0.57; Wilcoxon signed-rank test; n=244). Taken together, these findings confirm the hypothesis that the activity of transplanted cells is not required for the expression of OD plasticity, indicating rather that transplantation induces plasticity in the host circuit.

## Discussion

How plausible is that that a small number of transplanted interneurons, as few as 10% of the number of interneurons in the host tissue, could form a parallel circuit to directly inhibit the responses to the deprived eye? Because excitation and inhibition are finely balanced in the V1 cortical network (Troyer et al., 1998) an increase of the inhibition of the deprived eye’s pathway by 10% would be expected to reduce its response substantially, accounting for the OD shift. However, our findings reveal that transplanted neurons do not form a parallel circuit to mediate the expression of OD shift; rather they facilitate changes in the host circuit.

How do underlying neuronal mechanisms of transplant-induced CP compare to the normal CP in the postnatal visual cortex? The similarities can be recapitulated as follows. First, it has already been shown that the maturation of GABAergic inhibitory circuits is essential for the induction of the normal CP (Hensch, 2005; Espinosa and Stryker, 2012). Similarly, the timing of transplant-induced CP is dictated by the maturation of the transplanted cells (Southwell et al., 2010). Second, optogenetic suppression of host interneurons alters cortical responses but does not affect OD indices after MD during the normal CP (Saiepour et al., 2015), which indicates that the plasticity induced in the inhibitory connections does not selectively inhibit or disinhibit responses of deprived- or non-deprived-eye respectively. Here, our findings suggest a similar mechanism for transplant-induced OD plasticity. Our findings imply that OD plasticity in a property of the network and not merely of the inhibitory cells.

If the endogenous inhibitory circuit is compromised, the normal CP does not start (Fagiolini and Hensch, 2000). Likewise, transplanted interneurons that are unable to release GABA fail to induce a second CP (Priya et al., 2019). Specific changes in the activity of fast-spiking, parvalbumin-positive basket interneurons regulate the normal CP for OD plasticity (Kuhlman et al., 2011; Hengen et al., 2013; Figueroa Velez et al., 2017). However, either parvalbumin- or somatostatin-positive cells can induce a second CP after transplantation into the visual cortex (Tang et al., 2014). This indicates that MGE-derived interneurons appear to be unique in triggering subsequent competitive plasticity in excitatory connections (Larimer et al., 2016). However, CGE-derived interneurons are incompetent to induce OD plasticity.

Taken together, the present findings reveal that transplant-induced plasticity is not merely the result of a parallel circuit of transplanted cells but rather is a property of the host network that is induced but not maintained by the transplanted cells. How few numbers of transplanted neurons induce the second CP remains to be understood. One possible explanation consistent with theoretical studies of attractor networks (Aljadeff et al., 2015), is that inhibition provided by the transplanted cells perturbs the stability of the cortical network, allowing the relatively weak stimulus to change produced by monocular occlusion to push it out of its attractor state. The destabilized network could then rewire and settle into a new attractor state. Future work will be required to determine anatomical and functional changes in the synaptic connections.

## Acknowledgments

We thank members of the Stryker, Hasenstaub and Alvarez-Buylla laboratories for helpful discussion.

## Author contributions

M.S.H and M.P.S. designed research; M.S.H., B.R., and Q.F.R performed transplants; M.S.H. performed research and analyzed data; M.S.H. wrote the first draft of the paper; M.S.H., B.R., A.R.H., A.A.B., and M.P.S. edited the paper.

## Data and materials availability

All data, code, and materials used in the analysis are available online at https://github.com/Mahmood-Hoseini/TransplantRoleODIExpression-2019

## Ethics

Animal experimentation: This study was performed in strict accordance with the recommendations in the Guide for the Care and Use of Laboratory Animals of the National Institutes of Health. All of the animals were handled according to protocols approved by the UCSF institutional animal care and use committee under IACUC Protocol AN143347-03A.

